# Unravelling resistance mechanisms of oncolytic viruses in glioblastoma

**DOI:** 10.64898/2026.08.24.745745

**Authors:** Tine Deconinck, Tim Dierckx, Frederik De Smet, Jim Baggen, Dirk Daelemans

## Abstract

Glioblastoma (GBM) is an aggressive primary brain tumor with a major unmet medical need. Oncolytic viruses (OVs) show promise for GBM treatment, but complete remissions remain rare. The intratumoral heterogeneity of GBM drives therapeutic escape and emergence of OV-resistant subclones. Beyond the well-characterized interferon-mediated antiviral response, mechanisms driving OV resistance remain poorly understood. To identify new markers of tumor-intrinsic OV resistance in GBM, we exposed 14 GBM patient-derived cell lines (GBM-PDCLs) to 6 OVs and generated virus-resistant subpopulations from surviving cells. Focusing on Sindbis (SINV)- and H1-parvovirus (H1PV)-resistant cells, we showed that resistance is associated with impaired viral replication. Gene set enrichment analysis of transcriptomic profiles revealed that resistance to both SINV and H1PV correlated with downregulated glutamate receptor signaling. In contrast, collagen fibril organization was downregulated in SINV-resistant GBM PDCLs but upregulated in H1PV-resistant cells. Functional validation confirmed opposing effects of collagen degradation on SINV and H1PV oncolytic activity. One SINV-resistant GBM-PDCL showed cross-resistance to multiple OVs, which was associated with increased expression of antiviral immunity genes and increased dependence on type I interferon signaling for survival. Together, these findings reveal shared and virus-specific cellular processes driving OV resistance in GBM, providing a basis for strategies to overcome resistance.

## Introduction

Glioblastoma (GBM) is the most aggressive primary brain tumor in adults, with a median survival of approximately 14 months despite intensive multimodal treatment^1^. The current standard of care, maximal surgical resection followed by radiotherapy and temozolomide chemotherapy^2^, offers only temporary disease control, as nearly all patients experience tumor recurrence. This therapeutic failure is largely attributed to the high degree of intratumoral heterogeneity and the emergence of therapy-resistant tumor subclones^3^. Oncolytic viruses (OVs) have emerged as a promising alternative therapeutic approach for GBM. These viruses selectively infect and lyse malignant cells while sparing surrounding healthy brain tissue. The tumor specificity of OVs arises from various cancer-associated abnormalities, such as dysregulated interferon (IFN) or p53 signaling, hypoxic microenvironments, and elevated expression of viral entry receptors^4–6^. Beyond direct oncolysis, OVs can stimulate antitumor immunity by releasing tumor-associated antigens and danger-associated molecular patterns, thereby promoting immune-mediated tumor clearance^7^.

Most clinical trials involving OVs still rely on a ‘one-virus-fits-all’ approach, with limited consideration to patient stratification^8^. Resistance to OV therapy remains a key factor limiting efficacy and often leads to tumor recurrence. While antiviral immunity mediated through interferon (IFN) signaling is a well-recognized mechanism^9^, it likely represents only one facet of OV resistance. Non-immune tumor-intrinsic factors may also play a crucial role in the emergence of OV resistance and remain poorly understood. A more comprehensive understanding of the diverse mechanisms underlying OV resistance will be essential to move the field toward successful personalized virotherapy.

Several different experimental approaches have been applied for identification of tumor-intrinsic resistance factors of OVT for GBM. First, analysis of differential gene expression (DGE) between OV-responsive and non-responsive GBM cell lines is a common strategy to identify gene(s) (signatures) associated with OV resistance, which resulted in a prediction algorithm with the potential to preselect OVs or combinatorial strategies^10^. Second, CRISPR-Cas9 based genetic screening previously identified cellular factors involved in chromatin organization to be responsible for regulation of antiviral gene expression^11^. Third, our group previously conducted a comparative study in which the efficacy of multiple OVs was assessed across a large panel of GBM cell lines, followed by a correlation analysis to identify determinants of OV sensitivity and resistance^12^. That study demonstrated that viral-entry related molecules alone are generally insufficient for prediction of OV efficacy, with some notable exceptions. Importantly, 2 groups of OVs (hereafter referred to as “lysogroups”) were identified with opposing preferences for the GBM subtypes defined by Neftel *et al*.^13^ and with opposing associations with gene sets related to neurodevelopment, immune responses and extracellular matrix organization^12^.

These approaches primarily focus on baseline differences in gene expression profiles between responsive and non-responsive GBM tumors and therefore identify determinants of resistance present in the majority of tumor cells prior to treatment. Considerably less is known about less-pronounced, intrinsic mechanisms by which tumors can acquire resistance following OV treatment. This distinction is particularly relevant in GBM, where pronounced intratumoral heterogeneity can permit the survival of resistant subpopulations despite an initial response of the bulk tumor, ultimately leading to disease recurrence.

To investigate intrinsic resistance across distinct OV platforms, we selected Sindbis virus (SINV) and H1-parvovirus (H1PV). SINV is an enveloped, positive-sense RNA virus of the *Togaviridae* family that has been explored preclinically as an oncolytic agent for GBM, demonstrating safety and therapeutic efficacy when engineered to express cytokines^14^. In contrast, H1PV is a small, non-enveloped single-stranded DNA virus belonging to the *Parvoviridae* family that has shown a favorable safety profile and signs of immunogenic activity in a Phase I/IIa clinical trial in GBM patients^15^. These two viruses were selected because our previous comparative evaluation revealed differential sensitivity of GBM patient-derived cell lines (GBM-PDCLs) to SINV and H1PV, two OVs belonging to distinct lysogroups and characterized by different oncolytic mechanisms^12^, providing a unique opportunity to identify both shared and virus-specific mechanisms underlying intrinsic resistance to oncolytic virotherapy while maintaining experimental feasibility.

Here, we generated OV-resistant cell populations from GBM-PDCLs and compared gene expression profiles to identify those profiles associated with resistance to SINV and H1PV in GBM. Using this strategy, we identified downregulation of glutamate receptor signaling as a potential shared OV resistance mechanism, while degradation of collagen was associated with resistance to SINV and sensitivity to H1PV. Next to this, one of the SINV-resistant cell lines showed cross-resistance which was associated with an upregulation and acquired dependence on type I IFN signaling. Together, these results demonstrate that OV resistance can arise through both shared and virus-specific mechanisms, underscoring the need for biomarker-guided and adaptive therapeutic strategies to optimize the clinical efficacy of oncolytic virotherapy in GBM.

## Material and methods

### Cell lines

A total of 14 GBM-PDCLs were used in experiments reported in this manuscript. GBM-PDCLs, LBT and CME, were obtained from the Laboratory for Precision Cancer Medicine at KU Leuven. These PDCLs were established from fresh tumor tissue collected from patients undergoing surgical resection at UZ Leuven (all patients provided informed consent) or from retrospective dissociated tumor tissue samples obtained from the Center for Human Genetics. The Ethics committee Research UZ/KU Leuven approved the use of these established PDCLs under protocol S67312.

### Cell culture

GBM-PDCLs were cultured in NeuroCult™ NS-A Basal medium (human, StemCell technologies, #05750) with NeuroCult™ Proliferation Supplement (human, StemCell technologies, #05753), Human Recombinant basic Fibroblast Growth Factor (20 ng/mL, StemCell Technologies, #78003.2), Human Recombinant Epidermal Growth Factor (20 ng/mL, StemCell Technologies, #78006.2), 0.0002% Heparin sodium salt (StemCell Technologies, #07980), and 1% Antibiotic Antimycotic (Gibco, Thermo Fisher Scientific, #15240062). Plates or flasks were coated with 5 µg/ml laminin (Sigma-Aldrich, #L2020) at least 1 hour prior to seeding at 37°C. All cells were maintained at 37°C and 5% CO_2_ and were routinely tested for contamination by mycoplasma. Prior to passaging, cells were stained with Trypan Blue Stain (0.4%, Thermo Fisher Scientific, #15250) and counted with the LUNA-II Automated Cell Counter System (Logos Biosystems, #L40002).

### Viruses

In total, 7 different virus strains were investigated. Sindbis virus AR-339 strain (SINV; ATCC VR-1585), vaccinia virus Western Reserve strain (VV.WR; ATCC VR-1354) and measles virus Edmonton-Zagreb vaccine strain (MV.EZ; Serum Institute of India^16^), were propagated in VeroE6 cells (ATCC CRL-1586; RRID: CVCL_0574). H1 parvovirus Toolan strain (H1PV; ATCC VR356) was propagated in HeLa Chang Liver cells (Cytion; #300139; RRID: CVCL_0238). Propagation of SINV, VV.WR, MV.EZ and H1PV was performed in Dulbecco’s Modified Eagle Medium (DMEM, Thermo Fisher Scientific, #41965039). Vesicular stomatitis virus pseudotyped with the glycoprotein of the lymphocytic choriomeningitis virus^17^ (VSV.GP) was propagated in BHK21 clone 13 cells (RRID: CVCL_1915; virus and cells kindly gifted by Dr. Guido Wollmann from Medical University of Innsbruck), cultured in Glasgow’s MEM (GMEM, Thermo Fisher Scientific, #11710035) with 5% tryptose phosphate (Thermo Fisher Scientific, #18050039) and 1% Pen-Strep (Thermo Fisher Scientific, #15140122). Adenovirus type 5 with a deletion of 24 base pairs in the E1A region and a modification in the virus fiber with a, Arginyl-Glycyl-Aspartic acid(RGD)-4C motif (DNX-2401^18^; gifted by Prof. dr. Marta Alonso from Cima Universidad de Navarra) was propagated in A549 cells (ATCC CCL-185; RRID: CVCL_0023), cultured in Ham’s F12K medium (Thermo Fisher Scientific, #21127030). Virus stocks were produced by seeding their appropriate propagation cells in medium with 5-10% heat-inactivated FBS (HyClone) to reach a confluency of ~80% the next day. Then, medium was replaced by medium with 2.5-5% heat-inactivated FBS (HyClone) and cells were infected with virus at a multiplicity of infection (MOI) ranging from 0.01 to 0.1. When a cytopathogenic effect (CPE) was observed throughout the whole culture flask, supernatant was collected, centrifuged to remove cell debris, and stored at −80°C. After production, virus stocks were titrated by end-point dilution assays in the corresponding propagation cell line. Cells were infected with ten-fold serial dilutions of the virus for 7 days at 37°C, followed by visual scoring of CPE. The tissue culture infectious dose 50% (TCID50) was calculated using the method of Reed and Muench^19^. Allantoic fluid stocks of Newcastle disease virus La Sota strain (NDV.La.Sota; La Sota/Gallus_gallus/Belgium/634/2020) were purchased from Sciensano (Ixelles, Belgium).

### Generation of oncolytic virus resistant glioblastoma patient-derived cell lines

For generation of OV-resistant GBM-PDCLs, a panel of 14 GBM-PDCLs were selected with 6 different OVs: SINV, VSV.GP, VV.WR, H1PV and DNX-2401. GBM-PDCLs (50,000 cells/well) were seeded in a 24-well plate and infected with SINV, VSV.GP, VV.WR, H1PV or DNX-2401 at a MOI of 0.4, 0.2, 27.9, 17.8 or 0.6 respectively. For NDV La Sota, GBM-PDCLs were infected with 20µl of the purchased stock. Following infection, surviving cell populations were maintained in culture and expanded until sufficient cell numbers were obtained to perform resistance assessment using a cell viability assay. SINV- and H1PV-resistant subpopulations were subsequently selected for further characterization, including assessment of cross-resistance to H1PV, SINV, VSV.GP, DNX-2401, MV.EZ, VV.WR and NDV.La.Sota.

### Cell viability assay

To evaluate cell viability after viral infection, WT and resistant GBM-PDCLs (10,000 cells/well) were seeded in 96-well plates. The following day, cells were infected with virus dilutions ranging from 1/4 to 1/400,000,000. Seven days post-infection, medium was removed from the cells and replaced by 3-(4,5-dimethylthiazol-2-yl)-5-(3-carboxymethoxyphenyl)-2-(4-sulfophenyl)-2H-tetrazolium (MTS) reagent (CellTiter 96® AQueous Non-Radioactive Cell Proliferation Assay (MTS; Promega, #G5430)), 6-fold diluted in PBS. Absorbance at 490 nm was measured with the Spark^®^ Multimode Microplate Reader. Cell viability experiments were performed at least twice. For each experiment, we included 2 wells per cell-virus dilution condition. Relative viabilities were calculated based on the cell viability data with **formula (1)**:

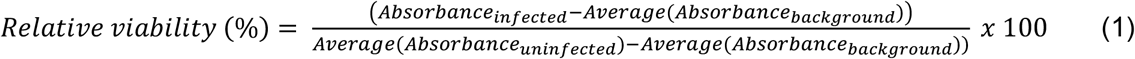

Multiplicity of infection (MOI) values were calculated for each virus dilution and dose-response curves based on the MOI (dose) and the relative viability (response) were made.

For experiments with collagenase I (#C1-BIOC, Sigma Aldrich), 5 hours prior to infection, cells were treated with 1 mg/ml collagenase I for 20 minutes. Seven days post-infection, MTS reagent was added directly to the medium in a 1:10 dilution.

### Compound testing

For experiments with ruxolitinib (INCB018424, #S1378, Sellekchem), GBM-PDCLs were seeded in 96well plates (10,000 cells/well) in 100 µL medium. The next day, a serial dilution of compound dissolved in 100 µL medium was added to the cells, with nine different concentrations as indicated in **Figure S3**. Seven days post-treatment, a cell viability assay was performed as previously described.

### Oncolytic score calculation

Area under the curve (AUC) values of dose-response curves were calculated using Graphpad Prism 10 (RRID:SCR_002798). The minimal relative viability corresponds to the lowest average relative viability obtained for a single MOI. As described before ^12^, an oncolytic score, combining the total AUC of the dose-response curve and the minimal relative viability that was reached, is used to quantify oncolytic efficacy of a virus and is calculated with **formula (2)**:

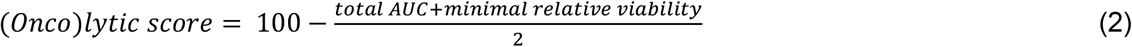

### Immunofluorescence staining

WT and resistant GBM-PDCLs were seeded at a density of 30,000 cells per well in 8-well µ-slides (ibidi). The following day, virus was added to the medium at MOI 0.04 for SINV, and MOI 1 for H1PV. After 48 hours (SINV) or 72 hours (H1PV), cells were fixed with 4% paraformaldehyde (PFA, Sigma-Aldrich, #252549), washed and permeabilized with 0.2% Triton X-100 (Sigma-Aldrich, #93443). Fixed cells were stained with mouse anti-dsRNA (J2; Scicons, #10010200, RRID:AB_2651015) at a 1:1000 dilution or undiluted anti-NS1 H1#742/1 monoclonal antibody, generated as described in Nüesch *et al*.^20^ (kindly gifted by Prof. dr. Jürg Nüesch from the German Cancer Research Center (DKFZ)). Secondary antibody Alexa Fluor® 488 goat anti-mouse (Thermo Fisher Scientific, #A11029) was diluted 1:500. Staining of collagen III was performed by fixing cells one day after seeding and staining with anti-collagen III (abcam, #ab184993) at a 1:100 dilution and secondary antibody Alexa Fluor®568 goat anti-rabbit at a 1:500 dilution. High-content imaging analysis (CellInsight CX5, Thermo Fisher Scientific) employing the HCS Studio Cell Analysis software (Thermo Fisher Scientific) was used for quantification of average pixel intensity per cell (NS1, collagen III) or percentage of cells with a fluorescent spot indicating replication (dsRNA). Staining experiments included two to four wells per condition and were performed twice.

### RT-qPCR

Total RNA was isolated from resistant and WT GBM-PDCLs using the RNeasy mini kit (#74104, Qiagen) and cDNA was synthesized with 500 ng of total RNA using the iScript cDNA synthesis kit (#1708890, BIO-RAD). RT-qPCR was performed on the resulting cDNA using the Sso Advanced universal SYBR green supermix (#1725274, BIO-RAD) together with the following primers: *COL1A1* forward (TCTGCGACAACGGCAAGGTG) and reverse (GACGCCGGTGGTTTCTTGGT) primers; *COL3A1* forward (GGAGCTGGCTACTTCTCGC) and reverse GGGAACATCCTCCTTCAACAG) primers. Reactions were run under the following conditions: 1 cycle at 50°C for 2 min and 95°C for 10 min, followed by 40 cycles at 95°C for 15 s and 60°C for 60 s. Real-time detection of the SYBR green fluorescence was conducted with QuantStudio™ 5 Real-Time PCR System (Applied Biosystems). For normalization, expression of the housekeeping gene *ACTB* was quantified for each sample using the following primers: *ACTB* forward (CCTTCCTGGGCATGGAGTCCTG) and reverse (GGAGCAATGATCTTGATCTTC). Fold-change values were calculated using the 2^−ΔΔCt^ method^21^.

### RNA sequencing and differential gene expression analysis

Total RNA of non-infected GBM-PDCLs CME016, LBT007/i and LBT123, established SINV-resistant LBT007/i and LBT123 and established H1PV-resistant CME016 and LBT123 GBM-PDCLs was extracted using the RNeasy kit (Qiagen, #74104). Four replicates of each sample were included and sequenced in one batch. RNA sequencing was performed by Novogene Europe (Cambridge, United Kingdom). Sequencing was performed on the Illumina Novaseq X platform and paired-end 150bp reads were generated. Reads containing adapters, N>10% (N represents unknown or uncalled bases) and low quality (Qscore≤5) base which is over 50% of the total base were removed. Trimmomatic (RRID:SCR_011848) was used to remove the first 15 bases from each read and to remove trailing bases with a Phred quality score below 3.

Read counts were generated by using Kallisto (RRID:SCR_016582; version 0.48.0)^22^ with 100 bootstraps and 8 threads. An index was generated using the GRCh38.96 transcriptome, downloaded from https://github.com/pachterlab/kallisto-transcriptome-indices/releases. Transcript-level abundance estimates were imported using tximport (RRID:SCR_016752; version 1.34.0)^23^ and summarized to gene level using the summarizeToGene function. Differential gene expression analysis was performed using DESeq2^24^ (RRID:SCR_015687; version 1.46.0). For each virus-specific analysis, samples were grouped according to the VirusResistance factor with two levels: WT and resistant, representing the parental GBM-PDCL and its corresponding virus-resistant derivative, respectively. Genes with zero counts in more than 3 samples were filtered out prior to analysis. The Wald test was used to assess differential expression, and p-values were adjusted using the Benjamini–Hochberg (BH) method. Log_2_ fold changes and adjusted p-values were calculated using the DESeq2 results function with the contrast vector c(“VirusResistance”, “virusX”, “WT”), where virusX represents the virus-specific resistant cell line and the resistant derivative is compared against its matched WT counterpart. Consequently, positive log_2_ fold change values indicate higher expression in resistant populations, whereas negative results indicate higher expression in WT cells.

To assess pathway-level changes in gene expression, we performed Gene Set Enrichment Analysis (GSEA) using the fgsea package^25^ (RRID:SCR_020938; version 1.32.2) with p-values calculated from 10,000 permutations. Gene sets, with a minimal size of 20, were obtained from the Molecular Signatures Database (MSigDB) C5 collection^26^, specifically from the GO Biological Process set.

### GBM subtype classification

GBM subtype scores were calculated as previously described^12^. Briefly, a Seurat (RRID:SCR_016341) object of the raw read counts of each GBM-PDCL was created and log-normalized. GBM subtype scores were calculated with the AddModuleScore function using the GBM meta-module gene lists from Neftel *et al*.^27^. The maximum subtype score determines the quadrant localization in **Figure S3A** and coordinates are determined based on log_2_(|SC_1_-SC_2_|+1) with SC_1_ and SC_2_ being the maximum subtype score and the subtype score of the neighboring quadrant, respectively. The GBM subtype plot was created using ggplot2 3.5.1 (RRID:SCR_014601).

### Generation of COL3A1 and COL1A1 knock-out cell lines

sgRNAs targeting COL3A1 (sg1: 5’-ACTCGCCCTCCTAATGGTCA-3’; sg2: 5’-AGGATGACCAGATGTACCAG-3’), COL1A1 (sg1: 5’-ATACTTACGACAGCGCCAGG-3’; sg2: 5’-CCAAGAAACCACCGGCGTCG-3’) and AAVS1 (5’-GTCACCAATCCTGTCCCTAG-3’) sgRNAs were cloned into the pU6-sgRNA plasmid using the Target Guide Sequence Cloning Protocol^28,29^. After transformation of NEB 5-alpha Competent E. coli (NEB, #C2987H), plasmid DNA was extracted with the Nucleospin Plasmid Purification Kit (Macherey-Nagel, #740588) and correct sgRNA integration was confirmed by Sanger sequencing.

Virus-like particles (VLPs) were generated in HEK293T cells via co-transfection of pBS-CMV-gagpol (RRID: Addgene_35614), pCMV-MMLVgag-3xNES-Cas9 (RRID: Addgene_181752), pCMV-VSV-G (RRID: Addgene_8454), and the pU6-sgRNA plasmid using an MLV-based production system^30^. HEK293T cells were plated in T150 flasks and transfected at 70–80% confluence using TurboFectin 8.0 (Origene, #TF81001). After 18 h, the medium was exchanged for DMEM supplemented with 1.1 g/100 mL BSA. VLP-containing supernatant was collected 48 h post transfection and clarified by centrifugation (200 × g, 3 min, 4 °C), taking care not to disturb cellular debris. VLPs were subsequently concentrated by overnight centrifugation at 4300 × g at 4 °C, resulting in visible white pellets. The following day, pellets were resuspended in 200 µL cold PBS containing 2% FBS, incubated for 1 h at 4 °C after which they were gently mixed, aliquoted, and stored at ™80 °C.

Cells were seeded (25,000 cells/well) in 24-well plates and 10 µL of the VLPs carrying Cas9 and each sgRNA were added to the culture medium. After 72 h, the VLP-containing medium was replaced with fresh growth medium, and cells were scaled up for subsequent use in the cell viability assay.

## Results

### Establishment of oncolytic virus-resistant patient-derived glioblastoma cell lines

To investigate resistance mechanisms to OV therapy in GBM, we aimed to derive virus-resistant subpopulations from GBM-PDCLs. We selected six OVs currently under (pre-)clinical evaluation for GBM treatment: Sindbis virus (SINV), glycoprotein-modified vesicular stomatitis virus (VSV.GP), vaccinia virus Western Reserve strain (VV.WR), adenovirus type 5 with a deletion in the E1A region and a modification in the virus fiber (DNX-2401), parvovirus H1 (H1PV), and Newcastle disease virus La Sota strain (NDV.La.Sota). A total of 14 GBM-PDCLs were infected with each virus and three distinct phenotypic outcomes were observed: a complete cytopathogenic effect (CPE), insensitivity or mild sensitivity to infection (weak selection), or initial strong sensitivity (stringent selection) with subsequent emergence of resistant clones (**Figure 1A**). The latter populations were expanded, yielding 18 surviving GBM subpopulations. To assess whether these populations remained virus-resistant, they were re-infected with a dilution series of the virus originally used in their selection. A population was considered resistant if its relative viability compared to the parental cell line was significantly (unpaired t test, n=2, p<0.05) increased for at least two virus dilutions, with an increase of ≥40% for at least one virus dilution. Seven subpopulations (39%) displayed sustained resistance (**Figure 1A and 1B)**, whereas resistance could not be confirmed in 11 subpopulations (61%) **(Figure S1**). In this way, we established two resistant cell lines for SINV, H1PV, and NDV.La.Sota and one resistant cell line for VSV.GP, while selection with VV.WR or DNX-2401 did not yield surviving populations. For subsequent analyses, we focused on SINV- and H1PV-resistant GBM-PDCLs.

**Figure 1:**
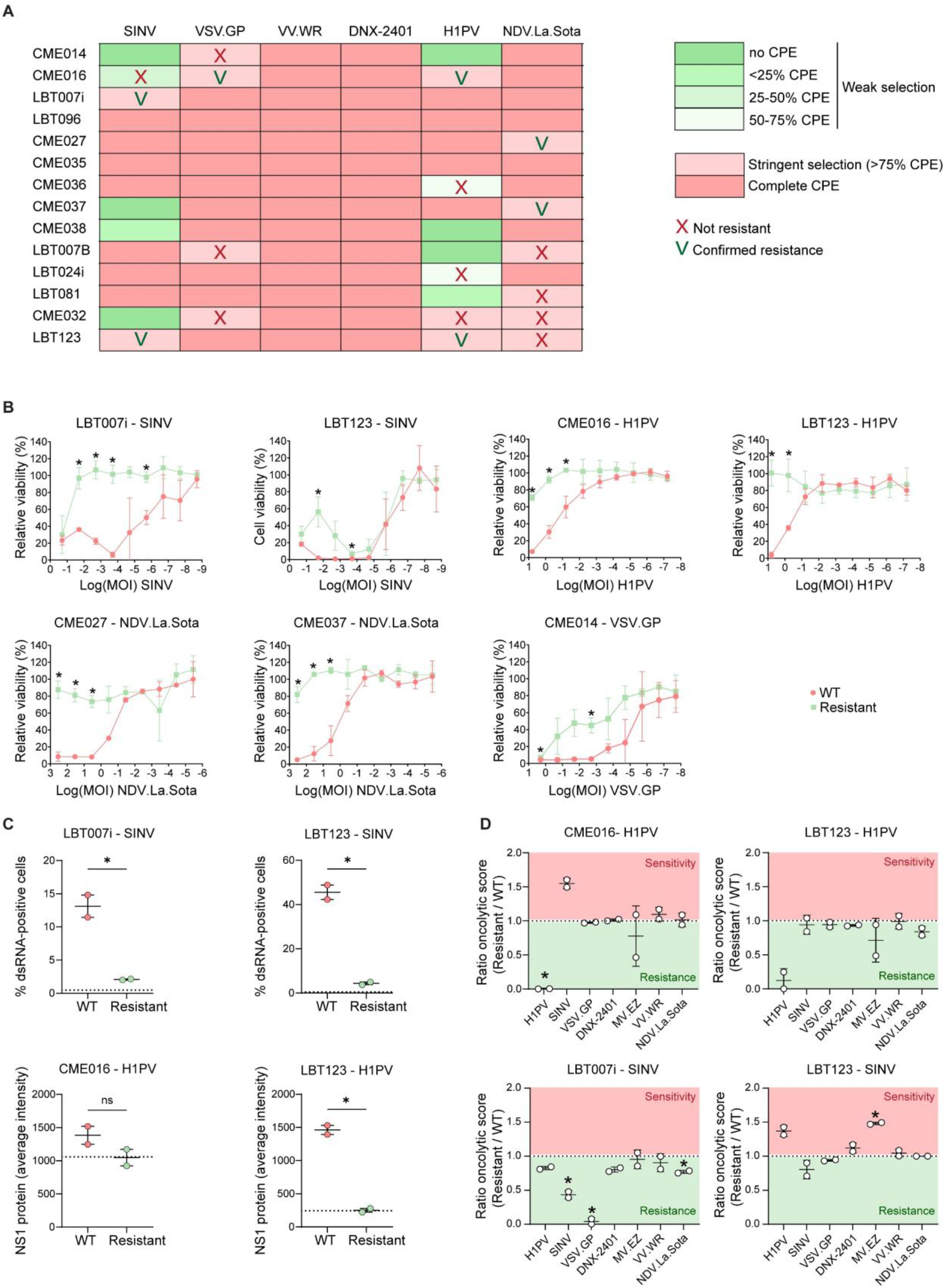
Establishment of oncolytic virus-resistant patient-derived glioblastoma cell lines with Sindbis- and H1-parvovirus-resistant cell lines showing impaired replication and virus-specific sensitivities. **(A)** Glioblastoma patient-derived cell line (GBM-PDCL) with a weak selection (less than 75% cytopathogenic effect (CPE)) are indicated in green, whereas stringent selection is indicated in pink. GBM-PDCLs that completely died after OV infection are indicated in dark pink. Cell lines that survived OV selection but did not show sustained resistance to the same virus are indicated with a red X, confirmed resistance is indicated with a green V. **(B)** Dose-response curves for each established OV-resistant GBM-PDCL. Viability, relative to uninfected control is shown on the y-axis, whereas the logarithmic function of the multiplicity of infection (MOI) of the virus is shown on the x-axis. An unpaired t test was used to compare the cell viabilities per MOI between resistant and wild type (WT) GBM-PDCLs. **(C)** Quantification of Sindbis (SINV) and H1-parvovirus (H1PV) replication as measured by immunofluorescence staining of dsRNA and NS1 protein, respectively. For H1PV, the average pixel intensity per cell is shown whereas for SINV, the percentage of cells with dsRNA staining is indicated on the y-axis. The dotted line represents the background signal measured in uninfected control cells. An unpaired t test was used to compare replication in resistant GBM-PDCL and WT counterpart. **(D)** Sensitivities of resistant versus WT GBM-PDCLs towards other OVs. The ratio of the oncolytic score in the resistant GBM-PDCL to the WT GBM-PDCL is shown on the y-axis. A value higher than 1 indicates more sensitivity to the OV in the resistant GBM-PDCL, while a value lower than 1 indicates more resistance in the resistant GBM-PDCL. A one sample t test was performed with test value 1. Mean of 2 experiments is shown and standard deviations are indicated for B-D. * = p<0.05.

### Impaired replication and virus-specific sensitivities in SINV- and H1PV-resistant glioblastoma patient-derived cell lines

To determine whether resistance to SINV and H1PV was associated with impaired viral replication, we performed immunofluorescence staining for viral replication markers. To quantify SINV replication, we immunostained dsRNA, an intermediate product of viral replication, whereas H1PV replication was quantified by staining non-structural protein 1 (NS1). We observed that in both SINV- and H1PV-resistant GBM-PDCLs, replication of the respective viruses was significantly (unpaired t test, n=2, p<0.05) decreased compared to WT cells (**Figure 1C**), except for the H1PV-resistant CME016 cell line. To identify potential cross-resistance to other OVs, resistant and WT GBM-PDCLs were infected with a serial dilution of either H1PV, SINV, VSV.GP, DNX-2401, MV.EZ, VV.WR or NDV.La.Sota. Oncolytic scores were calculated as previously described^12^ and the ratio of oncolytic scores relative to the WT was calculated for each virus in each pair of GBM-PDCLs (**Figure 1D**). As expected, the virus to which the GBM-PDCL subpopulation was confirmed to be resistant (**Figure 1B**) always displayed a ratio below 1, indicating a lower oncolytic score of this virus in resistant cells compared to their score in WT cells. In addition, the LBT007i SINV-resistant cell line showed significant cross-resistance towards other viruses: VSV.GP and NDV.La.Sota, while no cross-resistance was observed in the other resistant GBM-PDCLs. Notably, we observed reciprocal hypersensitivity: a ratio above 1 was detected for H1PV in SINV-resistant LBT123 cells and, conversely, for SINV in H1PV-resistant CME016 cells (**Figure 1D**). We previously showed that H1PV and SINV belong to different lysogroups, which have distinct oncolytic mechanisms and opposing preferences for GBM subtypes^12^. This might explain why a cell population can be resistant to one OV, while being hypersensitive towards another OV. To explore whether combining SINV and H1PV increases their cytolytic potential, we simultaneously infected parental CME016 cells with SINV and H1PV. We observed no marked difference between the single-virus and the combined conditions (**Figure S2**). Altogether, these results show that OV resistance is associated with impaired virus replication and that resistance to one OV can lead either to cross-resistance or to hypersensitivity toward other viruses. To gain deeper insight into the molecular basis of these distinct resistance phenotypes, we next performed gene expression analysis in the different GBM-PDCLs.

### Transcriptome profiling reveals cellular processes associated with both general and virus-specific resistance

To assess the mechanisms underlying OV resistance, we performed mRNA sequencing of parental and SINV/H1PV-resistant cell populations. Based on the expression profiles, the subtype scores were calculated as defined by Neftel *et al*.^13^. All resistant cell lines showed decreased astrocyte-like (AC), oligodendrocyte progenitor-like (OPC) and neural progenitor-like (NPC) scores compared to the parental cell line (**Figure S3A**). Only the H1PV-resistant LBT123 and SINV-resistant LBT007i lines exhibited a subtype switch, transitioning toward mesenchymal and NPC states, respectively, while all other resistant lines maintained the subtype classification of their WT counterparts (**Figure S3B**). To identify biological processes associated with resistance, the gene expression profiles were subjected to DGE analysis comparing parental and resistant populations. A significant normalized enrichment score (NES) indicates that genes from a given gene set are preferentially enriched at either the top (positive NES) or bottom (negative NES) of a gene list ranked by fold-change values from the DGE analysis. Examination of the 20 gene sets with the highest (top) or lowest (bottom) NES values in each parental versus resistant cell line comparison revealed extensive enrichment of biological processes associated with immune responses, neurodevelopment, and DNA replication (**Figure S4**). To identify the most important gene sets associated with resistance to each virus, we highlighted the 10 highest- and 10 lowest-ranked gene sets for SINV and H1PV, based on the median NES rank across the two resistant cell lines. For both viruses, resistance was associated with a downregulation of gene sets related to glutamate receptor signaling and neurodevelopment (**Figure 2A and 2B**), suggesting a common feature of viral resistance. Consistent with this finding, inhibition of the AMPA glutamate receptor has previously been shown to reduce SINV-mediated oncolysis in a patient-derived GBM cell line^12^. Furthermore, the ‘Neurogenesis’ gene set, which was previously linked to AC, NPC, and OPC GBM cell states by Neftel *et al*.,^13^ was significantly downregulated across all resistant GBM-PDCLs (**Table S1**), in agreement with the subtype score shifts observed above (**Figure S3A**). Notably, we found the gene set ‘Collagen Fibril Organization’ to be upregulated in H1PV-resistant cells, whereas the same gene set was downregulated in SINV-resistant cells (**Figure 2A, 2B** and **Figure S4**). Ranking of gene sets based on the difference in NES scores between SINV and H1PV revealed that multiple gene sets related to collagen formation and extracellular matrix (ECM) organization were present in the top 10 most divergent gene sets (**Figure 2C**). These findings suggest that ECM remodeling may reflect virus-specific adaptive responses, and may potentially underly the differential oncolytic activity of SINV and H1PV. Notably, these viruses were previously shown to belong to distinct lysogroups and to be opposingly associated with ECM-related biological processes^12^.

**Figure 2:**
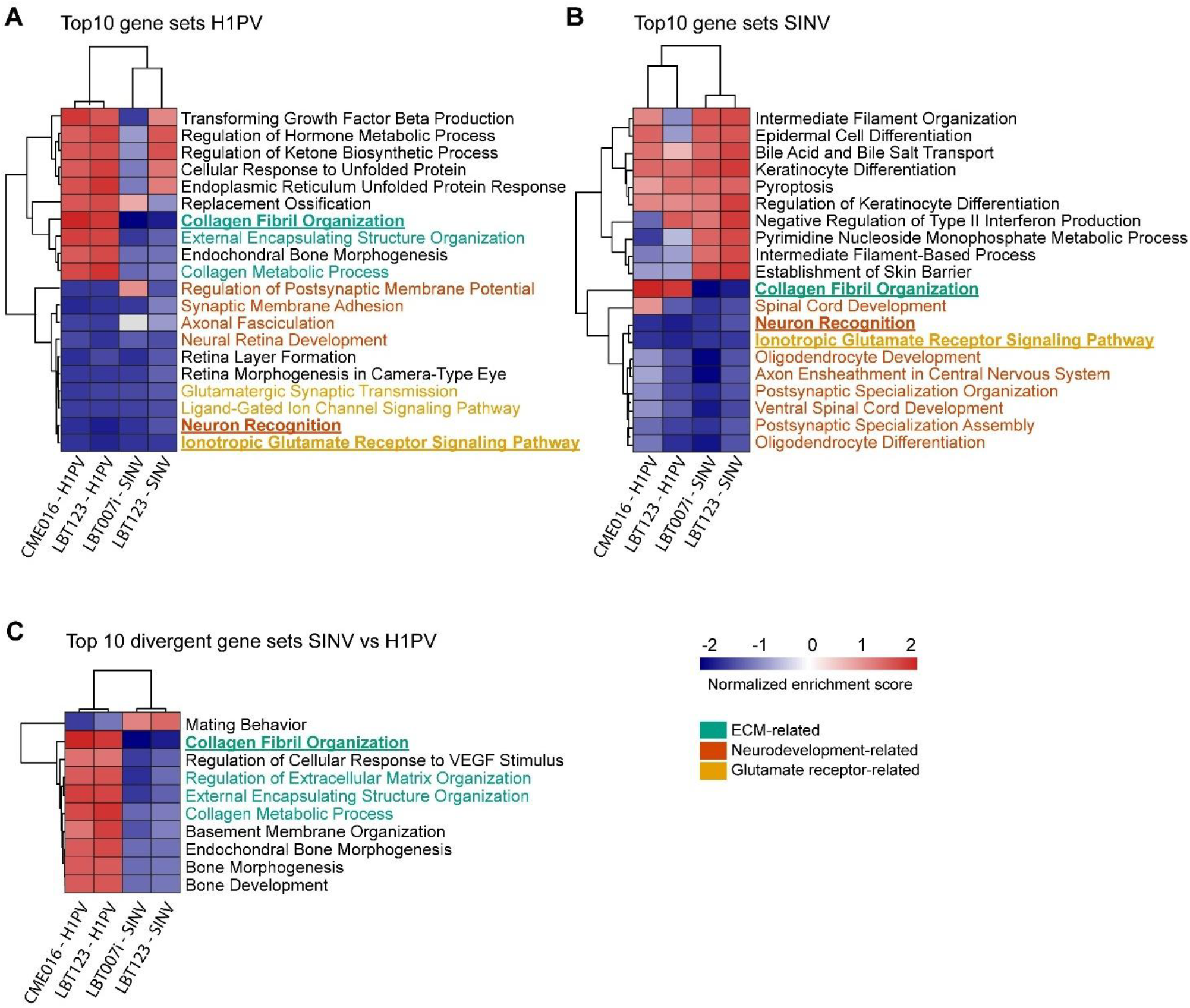
Transcriptome profiling reveals cellular processes associated with general and virus-specific resistance. Differential gene expression analysis between H1-parvovirus (H1PV)- and Sindbis virus (SINV)-resistant GBM-PDCLs and their WT counterpart was followed by gene set enrichment analysis. Median ranks of both SINV- or H1PV-resistant GBM-PDCLs were calculated based on their normalized enrichment scores. The gene sets with the 10 highest and lowest median ranks were selected for H1PV **(A)** and SINV **(B)** and the gene sets with the highest absolute difference in median ranks between H1PV- and SINV resistant GBM-PDCLs were identified **(C)**. Overlapping gene sets in the top 10 of H1PV and SINV are underlined and indicated in bold. Gene sets related to similar biological processes are indicated in green (extracellular matrix), dark orange (neurodevelopment) and yellow (glutamate receptor signaling). Hierarchical clustering is indicated for gene sets and GBM-PDCLs.

### Collagen degradation differentially affects oncolytic activity of SINV and H1PV

We next further explored the differential influence of collagen fibril organization on SINV and H1PV activity. Analysis of individual genes within the gene set ‘Collagen Fibril Organization’ revealed that the genes *COL1A1* and *COL3A1* have particularly strong opposite associations with SINV and H1PV resistance (**Figure 3A**). RT-qPCR analysis confirmed that these genes were upregulated in H1PV-resistant cells and downregulated in SINV-resistant cells (**Figure 3B**). At the protein level, immunofluorescence analysis revealed reduced collagen III levels in both SINV-resistant cell lines and elevated collagen III levels in the H1PV-resistant cell line LBT123 (**Figure 3C**), aligning with the RT-qPCR results. However, collagen III protein levels were reduced in the H1PV-resistant CME016 cell line, despite increased mRNA levels (**Figure 3B**). To determine whether ablation of these collagen-encoding genes influences sensitivity to H1PV or SINV, parental GBM-PDCLs were transduced with virus-like particles delivering two sgRNAs targeting either *COL3A1* or *COL1A1*, together with Cas9 protein to create genetic knock-out cells. Subsequent infection with either H1PV or SINV showed that knock-out of these individual genes did not significantly affect sensitivity to SINV or H1PV (**Figure S5**). To assess the broader role of collagens (**Figure 2A, 2B and Figure S4**) in mediating H1PV and SINV infection, we next examined whether general collagen degradation affects the oncolytic activity of either virus. To this end, the WT counterparts of the SINV- and H1PV-resistant GBM-PDCLs were pretreated with collagenase I, an enzyme that degrades triple-helical collagen fibrils, before infection with serial dilutions of each virus. Seven days post-infection, cell viability was quantified to evaluate oncolytic activity. Collagenase treatment induced detachment of GBM-PDCLs from the culture surface and promoted the formation of spheroid-like structures. In LBT123 and CME016 cells, pre-treatment with collagenase I significantly increased the oncolytic activity of H1PV at least for one MOI (**Figure 3D**). In contrast, collagenase I pre-treatment significantly reduced the oncolytic activity of SINV in LBT123 and LBT007i for at least 2 MOIs (**Figure 3E**). Together, these findings indicate that global collagen degradation, rather than specific targeting of *COL3A1* or *COL1A1*, facilitates H1PV-mediated oncolysis, whereas it impairs SINV-mediated oncolysis in GBM-PDCLs.

**Figure 3:**
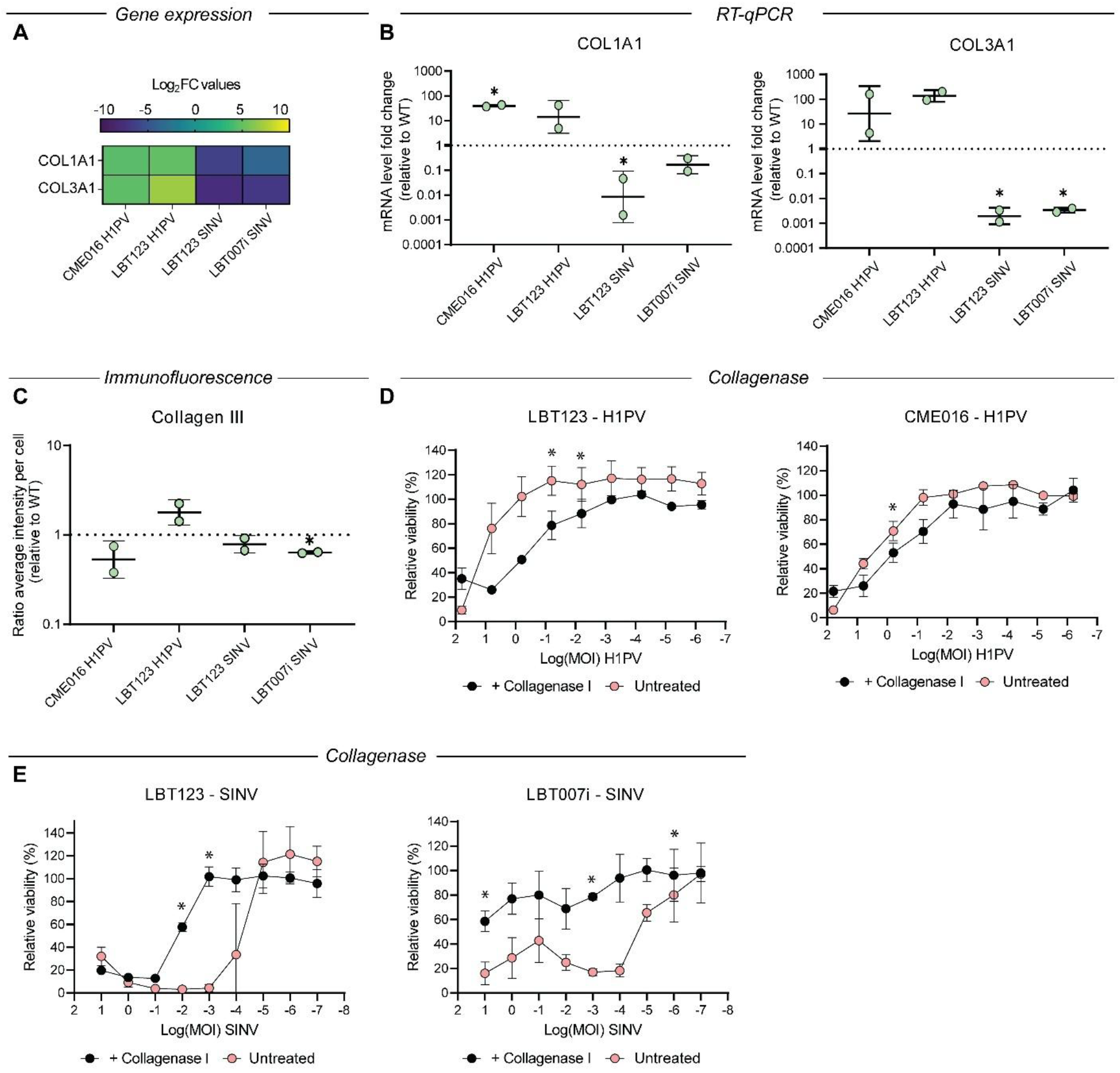
Collagen degradation differentially affects oncolytic activity of Sindbis and H1-parvovirus. **(A)** Log_2_FC values of the collagen fibril organization driver genes for both H1-parvovirus (H1PV)- and Sindbis virus (SINV)-resistant glioblastoma patient-derived cell lines (GBM-PDCLs). **(B)** RT-qPCR results for COL1A1 and COL3A1 in H1PV- and SINV-resistant GBM-PDCLs, compared to the WT GBM-PDCL. Relative expression was calculated using the 2^−ΔΔCt^ method, with C_t_ values for ACTB as reference. Unpaired t tests were performed for comparison between WT and resistant GBM-PDCL conditions. **(C)** Quantification of immunofluorescence staining using an anti-collagen III antibody in H1PV- and SINV-resistant GBM-PDCLs. Unpaired t tests were performed for comparison between WT and resistant GBM-PDCL conditions. **(D-E)** Viability, relative to uninfected control is shown for each MOI of H1PV **(D)** or SINV **(E)**, with and without pre-treatment with collagenase I. Paired t tests were performed for comparison between conditions with and without collagenase I treatment. Mean of 2 experiments is shown and standard deviations are indicated for B-E.*=p<0.05.

### Transcriptomic profiling reveals upregulation of interferon-mediated antiviral signaling underlying oncolytic virus cross-resistance

In contrast to the other virus-resistant cell lines, SINV-resistant LBT007i showed significant cross-resistance towards VSV.GP and NDV.La.Sota and potentially weak resistance to H1PV and DNX-2401 (**Figure 1D**). To explain this cross-resistance phenotype, the 20 most enriched and depleted gene sets identified in this LBT007i SINV-resistant subpopulation were compared across all resistant cell lines (**Figure 4A**). Gene sets related to regulation of viral processes and response to type I interferon were among the most strongly enriched pathways in SINV-resistant LBT007i, whereas they showed little or no enrichment in the other resistant subpopulations. These findings suggest that transcriptional activation of antiviral defense programs, particularly type I interferon signaling, may contribute to the unique cross-resistance phenotype observed in SINV-resistant LBT007i cells. As type I interferon signaling is mediated through JAK-STAT pathway, we hypothesized that SINV-resistant LBT007i cells had become increasingly dependent on this signaling axis and would therefore display increased sensitivity to JAK inhibition. To evaluate this possibility, we compared the sensitivity of WT and SINV-resistant LBT007i cells to ruxolitinib, a clinically approved JAK1/2 inhibitor that blocks downstream interferon signaling. SINV-resistant LBT007i cells displayed a 29-fold increase in ruxolitinib sensitivity compared with their WT counterpart (**Figure 4B**). These findings are consistent with enhanced reliance on IFN-I/JAK-STAT signaling in SINV-resistant LBT007i cells and suggest that acquisition of OV resistance may create a collateral vulnerability to JAK-STAT inhibition. Altogether, these results reveal a potential opportunity to counter OV resistance via sequential therapeutic targeting of JAK1 in OV-resistant tumor cell populations, rather than concurrent antiviral pathway suppression.

**Figure 4:**
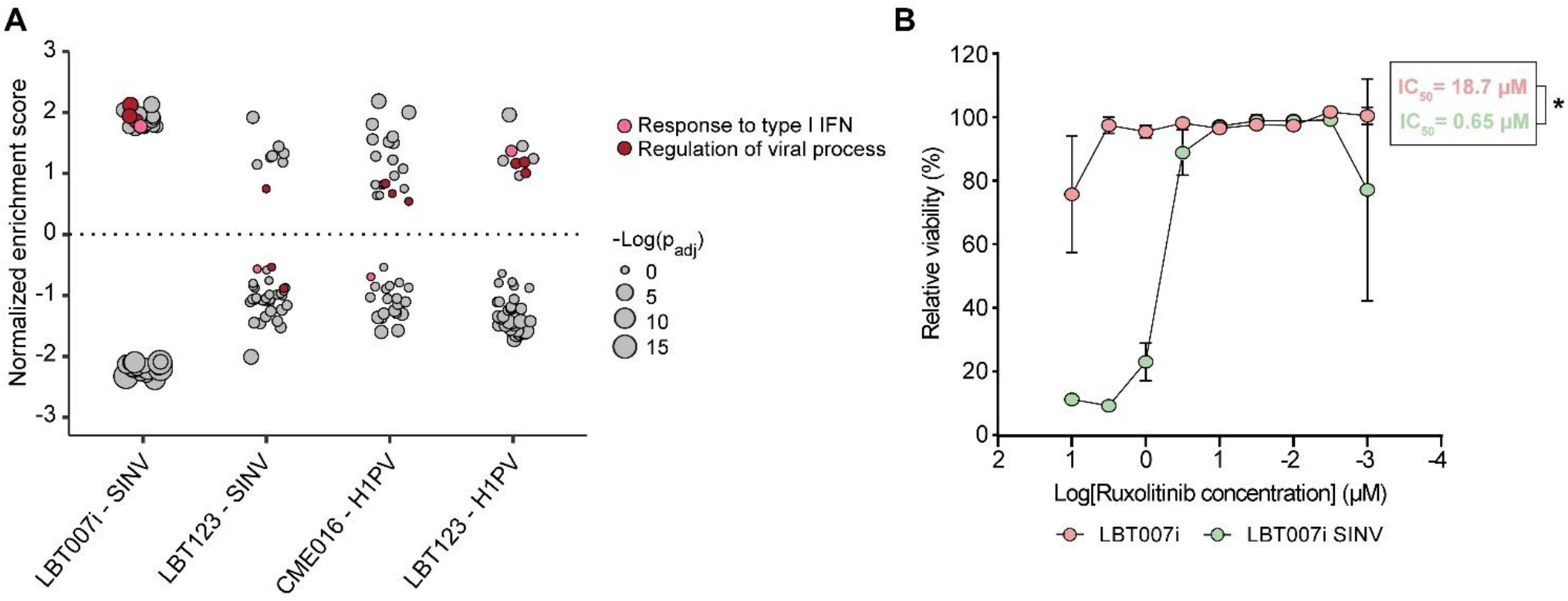
Transcriptomic profiling reveals upregulation of interferon-mediated antiviral signaling underlying oncolytic virus cross-resistance. **(A)** Normalized enrichment scores of the top 20 (enriched) and bottom 20 (depleted) gene sets of Sindbis (SINV)-resistant LBT007i relative to parental LBT007i cells are shown for all resistant cell lines. Dot size corresponds to the statistical significance of enrichment (negative logarithmic function of the adjusted p value). The gene set ‘Response to type I interferon’ is highlighted in pink, whereas gene sets related to regulation of viral processes are highlighted in red. **(B)** Viability, relative to uninfected control is shown on the y-axis, whereas the logarithmic function of the concentration of ruxolitinib (in µM) is shown on the x-axis. Nonlinear regression analysis revealed a significant difference in IC_50_ values between LBT007i and LBT007i SINV cells (extra sum-of-squares F test, F(1,32) = 125.4, p < 0.0001).

## Discussion

Resistance to OVs remains a major challenge for the effective treatment of GBM, yet the underlying mechanisms are incompletely understood. In this study, we generated OV-resistant derivatives of GBM-PDCLs and established stable resistance to four distinct OVs: SINV, H1PV, VSV.GP, and NDV.La.Sota. Focusing on SINV- and H1PV-resistant subpopulations, we found that resistance was consistently associated with reduced viral replication and a shared transcriptional program characterized by downregulation of glutamate receptor signaling and decreased expression of AC-, OPC-, and NPC-like cellular states. At the same time, our analyses uncovered virus-specific resistance mechanisms, identifying collagen fibril organization as a determinant with opposing effects on SINV and H1PV susceptibility. Functional experiments confirmed this observation, demonstrating that collagen degradation enhances H1PV-mediated oncolysis while impairing SINV-mediated oncolysis. Finally, we identified SINV-resistant LBT007i as a particularly distinct resistant model that exhibited broad cross-resistance to multiple OV platforms and relied on chronic type I IFN signaling for survival, highlighting potential vulnerabilities that may be exploited to overcome resistance.

In most surviving GBM-PDCL subpopulations (11/18; 61,1%), there was no sustained resistance (**Figure 1A**). This may be explained by the use of a single round of selection, which may permit the survival of incompletely infected cells or cells adopting transient, reversible stress responses rather than establishing stable resistance programs. Although additional rounds of selection may have yielded more stable resistant populations, their implementation was limited by the slow proliferation of GBM PDCLs, their strong dependence on cell-cell interactions, and the extended recovery period required following viral infection. Furthermore, SINV-resistant CME016 and H1PV-resistant CME036 and LBT024i were exposed to relatively weak selective pressure, as infection induced only 25-50% or 50-75% CPE. Notably, no surviving populations emerged after infection with VV.WR or DNX 2401 (**Figure 1A**), suggesting either that resistance to these viruses is less readily acquired under *in vitro* conditions or that the selective pressure applied was too stringent. Alternatively, resistance mechanisms relevant to VV.WR or DNX 2401 may depend on features of the *in vivo* tumor microenvironment, such as stromal support, immune interactions, spatial heterogeneity, or incomplete viral penetration, none of which are adequately recapitulated in conventional cell culture systems.

We prioritized SINV and H1PV resistant GBM PDCLs for mechanistic characterization. The consistent downregulation of neurodevelopmental-related gene sets across all resistant subpopulations was accompanied by reduced AC, OPC and NPC-like subtype scores (**Figure 2A, 2B and Figure S3A**). However, these transcriptional changes did not translate into a general shift in dominant cellular state (~maximal subtype) as most SINV- or H1PV-resistant GBM-PDCLs retained their parental subtype identity (**Figure S3B**). Instead, resistance seemed to be more closely associated with decreased expression of genes involved in glutamate receptor signaling. This observation is in line with our previous findings showing that pharmacological inhibition of the AMPA glutamate receptor reduces oncolytic activity of SINV, while H1PV-mediated oncolysis remains unaffected.^12^ Notably, glutamate signaling has also been implicated in SINV-induced neuronal death through excitotoxic mechanisms^31^, raising the possibility that this pathway contributes directly to virus-mediated tumor cell killing. Suppression of glutamate receptor signaling may attenuate OV-induced cytotoxicity and thereby promote resistance. While these findings suggest that enhancing glutamate signaling could potentially sensitize GBM tumors to OV therapy, such an approach would require careful consideration. Glutamate signaling is a well-established driver of glioblastoma proliferation, invasion, and resistance, and increased pathway activity has been linked to reduced responses to several clinically approved GBM treatments^32–34^. Therefore, any strategy aimed at exploiting glutamate signaling to enhance OV efficacy would need to balance potential gains in viral oncolysis against the risk of promoting tumor progression.

A key finding in this study is the identification of ‘Collagen Fibril Organization’ as the pathway most strongly and differentially associated with OV resistance in a virus-dependent manner, being upregulated in H1PV-resistant cells and downregulated in SINV-resistant cells (**Figure 2A, 2B and 2C**). Importantly, our functional data suggest that this association cannot be attributed to changes in a small number of individual genes. Collagen biology is inherently network-driven, requiring coordinated regulation of multiple genes to produce meaningful alterations in ECM composition and structure^35^. In line with this concept, knock-out of individual collagen genes (e.g., *COL1A1, COL3A1*) had only limited effects on viral sensitivity (**Figure S5**). In contrast, broad enzymatic degradation of collagen fibrils using collagenase induced clear and opposing phenotypes, enhancing H1PV-mediated oncolysis while reducing SINV-mediated oncolysis (**Figure 3D and 3E**). These findings indicated that the overall collagen matrix, rather than single collagen components, influences cellular susceptibility to OV infection and killing. Interestingly, *COL3A1* mRNA expression was increased in H1PV-resistant CME016 cells relative to their parental counterpart, while collagen III protein levels were decreased. Given that collagen III is a homotrimer composed of three identical α1(III) chains encoded by *COL3A1*, this discrepancy suggests regulation beyond transcription, potentially involving impaired translation, defective post-translational processing and fibril assembly, or increased protein turnover. Together, these observations highlight the importance of ECM remodeling in OV resistance and reveal fundamentally distinct interactions between collagen-rich microenvironments and the oncolytic activity of SINV and H1PV.

In a previous study^12^, we identified an inverse relationship between the oncolytic activities of H1PV and SINV, two viruses belonging to distinct lysogroups, and the expression of the MSigDB C5 Biological Process gene sets ‘Extracellular Structure Organization’ and ‘Extracellular Matrix Component’. These findings suggested that ECM remodeling may differentially affect the oncolytic activity of these viruses. The results presented here further support this hypothesis, as collagen fibril organization emerged as a resistance-associated pathway displaying opposite patterns in H1PV- and SINV-resistant cells. Together, the data indicate that ECM remodeling is not a universal determinant of susceptibility to OVs but rather exerts virus-specific effects that likely depend on the biological characteristics and infection strategies of individual viral lysogroups. These results highlight the importance of considering tumor ECM composition when evaluating or optimizing OV-based therapies.

Our results with H1PV are consistent with previous studies demonstrating that ECM components can act as both physical and biochemical barriers to restrict OV spread within solid tissues^36^. For example, an oncolytic herpesvirus type 1 engineered to express a matrix metalloproteinase showed improved intratumoral penetration and enhanced antitumor efficacy in a GBM xenograft mouse model, highlighting the therapeutic potential of ECM targeting strategies to facilitate viral spread^37^. In this context, approaches aimed at remodeling or degrading the ECM to enhance OV delivery and distribution may represent a promising avenue for enhancing H1PV oncolytic efficacy. However, our observations with SINV demonstrate that the benefits of ECM modulation are not universal and my vary substantially between OV platforms. Indeed, collagen degradation reduced SINV-mediated oncolysis, indicating that ECM-targeted interventions could be detrimental in certain virus-tumor contexts. These findings underscore the importance of evaluating ECM-modulating strategies on a virus-specific basis rather than assuming a broadly beneficial effect across OV therapies. More generally, our results align with increasing reports of the importance of ECM composition and architecture as critical determinants of GBM biology. Collagen-associated genes such as COL3A1^38^, have been implicated in tumor invasion, angiogenesis, and therapeutic resistance, while studies employing engineered ECM/3D systems have shown that matrix parameters can reshape tumor phenotypes^39,40^. Collectively, these observations support the notion that ECM remodeling is both a key regulator of tumor biology and an important determinant of OV responsiveness, warranting further investigation as context-dependent therapeutic target.

Among our established SINV- and H1PV-resistant GBM-PDCLs, SINV-resistant LBT007i was particularly notable because of its pronounced cross-resistance to multiple OVs (**Figure 1D**), suggesting engagement of a shared antiviral program rather than virus-specific escape mechanisms. Consistent with this, resistance was associated with a sustained upregulation of type I IFN response genes (**Figure 4A and Figure S4**). Interestingly, SINV-resistant LBT007i cells also displayed increased sensitivity to JAK/STAT inhibition by ruxolitinib (**Figure 4B**), suggesting a functional dependence on chronic type I IFN signaling. These findings are consistent with previous reports identifying innate antiviral signaling as a central barrier to OV efficacy, including resistance to reovirus^41^, HSV-1^42^, MV^43^, and VSV^44^. Together, these findings suggest that OV-resistant tumors with sustained type I IFN pathway activation may harbor a therapeutically exploitable vulnerability to JAK1/2 inhibition. Pharmacological targeting with ruxolitinib could therefore represent a rational strategy, although further validation *in vivo* is required.

Taken together, our data highlight the complexity and multifactorial nature of OV resistance in GBM, involving broad antiviral programs such as the type I IFN response, cell-state remodeling characterized by reduced AC, OPC, NPC populations, and virus-specific determinants including collagen organization. These findings argue against a “one-virus-fits-all” approach and suggest that OV monotherapy may be insufficient to achieve durable tumor control. Instead, multimodal or multi-virus strategies could exploit complementary mechanisms of action, whereby resistance to one virus is accompanied with increased susceptibility to another. This concept is supported by our observation that SINV-resistant cells exhibited enhanced sensitivity to H1PV, whereas H1PV-resistant cells became more sensitive to SINV, respectively (**Figure 1D**). In contrast, simultaneous SINV/H1PV co-infection did not provide additional benefit (**Figure S2**), suggesting that sequential rather than simultaneous OV administration may represent a more effective therapeutic strategy. As SINV and H1PV belong to distinct lysogroups^12^ with different mechanisms of action and preferential activity across GBM subtypes, determining the lysogroup sensitivity profile of a patient’s tumor may help identify the most suitable OV platform and predict opportunities for sequential multi-OV therapy. Such an approach could facilitate the rational selection of complementary viruses, thereby minimizing cross-resistance and maximizing therapeutic efficacy. Remarkably, a GBM patient who received a combination of multiple OVs experienced exceptional long-term disease control, remaining progression-free for 14.5 years following the initiation of therapy^45^. Although based on an isolated clinical observation, this case report further illustrates the promise of exploiting complementary multi-OV strategies for GBM to broaden tumor coverage and overcome intratumoral heterogeneity. Additionally, rational combination therapies, such as transient JAK/STAT inhibition or modulation of collagen remodeling, may help overcome resistance in specific molecular contexts. Collectively, these findings underscore the need for systematic characterization of GBM tumors to predict and therapeutically address both intrinsic and acquired mechanisms of OV resistance.

An important limitation of this study is that all experiments were conducted *in vitro* and therefore do not recapitulate the full complexity of the tumor microenvironment, including immune components, stromal interactions, and ECM architecture. These factors are known to critically influence OV efficacy and therapeutic response. Consequently, *in vivo* studies will be required to confirm our findings and further elucidate the underlying resistance mechanisms. Nevertheless, a major strength of this work lies in the use of multiple patient-derived GBM cultures, which more accurately capture the inter-/intra-tumoral heterogeneity and clinically relevant cell states observed in human tumors than conventional cell lines^46^. Moreover, the parallel analysis of SINV- and H1PV-resistant subpopulations enabled the distinction between shared and virus-specific resistance mechanisms.

In conclusion, we identify both common and virus-specific resistance programs that limit the efficacy of oncolytic virotherapy. While activation of innate antiviral signaling pathways and

shifts in neurodevelopmental cell-state emerge as shared hallmarks of resistance, collagen fibril organization exerts opposing effects on susceptibility to SINV and H1PV. These insights highlight the importance of comprehensive tumor characterization prior to OV therapy and support the development of adaptive treatment strategies tailored to the resistance landscape of individual tumors. Strategies such as sequential OV administration or rational combination therapies may help overcome resistance, enhance treatment efficacy, and ultimately improve clinical outcomes.

## Data availability statement

Data reported in this study will be shared by the corresponding author upon request. Raw read counts of WT, SINV-and H1PV-resistant GBM-PDCLs are available in **Table S2**.

## Acknowledgements

We thank Prof. dr. Marta Alonso from the University Clinic of Navarra for generously providing DNX-2401 and Dr. med. Guido Wollmann from the Medical University Innsbruck for generously providing VSV.GP used in this study. We thank Leentje Persoons and Prof. Graciela Andrei for their help with the collection of the viruses used in this study. We thank Marleen Derweduwe and Annelies Claeys from the Laboratory of Precision Medicine for their input on the origin and culturing of the GBM-PDCLs used in this study and their help with the RNA sequencing and application of the project to the Ethical Committee Research UZ/KU Leuven. We thank Brigitte Bais for her help with the establishment of the resistant cell lines, the optimalisation of the immunofluorescent staining and the preparation of samples for RNA sequencing. We thank Joni Punjwani, Yanicka Smolders, Nathalie Van Winkel and Niels Willems for their help with the immunofluorescent staining and Els Vanstreels for her help with the analysis of the immunofluorescent staining. This research was funded by the Research Foundation – Flanders (Fonds Wetenschappelijk Onderzoek, FWO; project number: 11M2425N) and the KU Leuven Special Research Fund (Bijzonder Onderzoeksfonds, BOF; project number: 3M240253). The Large Language Model Copilot 3.3.11 was used to improve the quality of the writing for specific sections of the manuscript.

## Author contributions

J.B., D.D., T. Deconinck and T. Dierckx contributed to the conception and design of the study. T. Deconinck conducted all experiments in this study. J.B., T. Deconinck and T. Dierckx were involved in the data processing and analysis. T. Deconinck wrote the first draft of the manuscript. J.B., D.D. and T. Dierckx contributed to the writing of the manuscript. D.D. supervised and supported the project. F.D.S. provided the GBM-PDCLs and supervised the project.

## Competing interests

The authors declare no competing interests

## Notes

### Competing Interest Statement

The authors have declared no competing interest.

## References

1. Ostrom, Q. T. et al. The epidemiology of glioma in adults: A state of the science review. Neuro. Oncol. 16, 896–913 (2014).

2. Stupp, R. et al. Radiotherapy plus concomitant and adjuvant temozolomide for glioblastoma. New England Journal of Medicine 352, 987–96 (2005).

3. De Vleeschouwer, S. (Ed.) Glioblastoma. Codon Publications (2017). doi:10.15586/codon.glioblastoma.

4. Howells, A., Marelli, G., Lemoine, N. R. & Wang, Y. Oncolytic viruses-interaction of virus and tumor cells in the battle to eliminate cancer. Front. Oncol. 7, (2017).

5. Kim, M. et al. The viral tropism of two distinct oncolytic viruses, reovirus and myxoma virus, is modulated by cellular tumor suppressor gene status. Oncogene 29, 3990–3996 (2010).

6. Jhawar, S. R. et al. Oncolytic viruses-natural and genetically engineered cancer immunotherapies. Front. Oncol. 7, (2017).

7. Marelli, G., Howells, A., Lemoine, N. R. & Wang, Y. Oncolytic viral therapy and the immune system: A double-edged sword against cancer. Front. Immunol. 9, (2018).

8. Stavrakaki, E., Dirven, C. M. F. & Lamfers, M. L. M. Personalizing oncolytic virotherapy for glioblastoma: In search of biomarkers for response. Cancers (Basel). 13, 1–22 (2021).

9. Bhatt, D. K. & Chammas, R. Resistance Mechanisms Influencing Oncolytic Virotherapy, a Systematic Analysis. Vaccines (Basel). 9, (2021).

10. Kurokawa, C. et al. Constitutive Interferon Pathway Activation in Tumors as an Efficacy Determinant Following Oncolytic Virotherapy. J Natl Cancer Inst 110, 1123–1132 (2018).

11. Guo, C. et al. BRD9 inhibition overcomes oncolytic virus therapy resistance in glioblastoma. Cell Rep. Med. 6, 102258 (2025).

12. Deconinck, T., Dierckx, T., De Smet, F., Baggen, J. & Daelemans, D. Comparative evaluation of oncolytic viruses reveals opposing preferences for glioblastoma subtypes. Molecular Therapy Oncology 34, 201263 (2026).

13. Neftel, C. et al. An Integrative Model of Cellular States, Plasticity, and Genetics for Glioblastoma. Cell 178, 835–849.e21 (2019).

14. Sun, K. et al. Oncolytic Viral Therapy for Glioma by Recombinant Sindbis Virus. Cancers (Basel). 15, 1–15 (2023).

15. Geletneky, K. et al. Oncolytic H-1 Parvovirus Shows Safety and Signs of Immunogenic Activity in a First Phase I/IIa Glioblastoma Trial. Molecular Therapy 25, 2620–2634 (2017).

16. Serum Institute of India. MEASLES VACCINE, LIVE, ATTENUATED (FREEZE-DRIED). https://www.seruminstitute.com/product_viral_measles.php.

17. Muik, A. et al. Pseudotyping Vesicular Stomatitis Virus with Lymphocytic Choriomeningitis Virus Glycoproteins Enhances Infectivity for Glioma Cells and Minimizes Neurotropism. J. Virol. 85, 5679–5684 (2011).

18. Lamfers, M. L. M. et al. Potential of the conditionally replicative adenovirus Ad5-Delta24RGD in the treatment of malignant gliomas and its enhanced effect with radiotherapy. Cancer Res. 62, 5736–42 (2002).

19. Reed, L. j. & Muench, H. A Simple Method Of Estimating Fifty per Cent Endpoints. Am. J. Epidemiol. 27, 493–497 (1938).

20. Tessmer, C. et al. Generation and Validation of Monoclonal Antibodies Suitable for Detecting and Monitoring Parvovirus Infections. Pathogens 11, 1–20 (2022).

21. Livak, K. J. & Schmittgen, T. D. Analysis of relative gene expression data using real-time quantitative PCR and the 2-ΔΔCT method. Methods 25, 402–408 (2001).

22. Bray, N. L., Pimentel, H., Melsted, P. & Pachter, L. Near-optimal probabilistic RNA-seq quantification. Nat. Biotechnol. 34, 525–527 (2016).

23. Soneson, C., Love, M. I. & Robinson, M. D. Differential analyses for RNA-seq: transcript-level estimates improve gene-level inferences. F1000Res. 4, 1521 (2015).

24. Love, M. I., Huber, W. & Anders, S. Moderated estimation of fold change and dispersion for RNA-seq data with DESeq2. Genome Biol. 15, 1–21 (2014).

25. Korotkevich, G., Sukhov, V. & Sergushichev, A. Fast gene set enrichment analysis (R package). (2021).

26. Liberzon, A. et al. Molecular signatures database (MSigDB) 3.0. Bioinformatics 27, 1739–1740 (2011).

27. Neftel, C. et al. An Integrative Model of Cellular States, Plasticity, and Genetics for Glioblastoma. Cell 178, 835–849.e21 (2019).

28. Sanjana, N. E., Shalem, O. & Zhang, F. Improved vectors and genome-wide libraries for CRISPR screening. Nat Methods 11, 783–784 (2014).

29. Shalem, O. et al. Genome-Scale CRISPR-Cas9 Knockout. Science (1979). 343, (2014).

30. Banskota, S. et al. Article Engineered virus-like particles for efficient in vivo delivery of therapeutic proteins ll ll Engineered virus-like particles for efficient in vivo delivery of therapeutic proteins. Cell 185, 250–265 (2022).

31. Nargi-Aizenman, J. L. & Griffin, D. E. Sindbis Virus-Induced Neuronal Death Is both Necrotic and Apoptotic and Is Ameliorated by N-Methyl-d-Aspartate Receptor Antagonists. J. Virol. 75, 7114–7121 (2001).

32. Radin, D. P. AMPA Receptor Modulation in the Treatment of High-Grade Glioma: Translating Good Science into Better Outcomes. Pharmaceuticals 18, 33–38 (2025).

33. Wilkins, D. & Kumar, S. Glutamate Drives Glioblastoma Invasion in Three-Dimensional Hyaluronic Acid Hydrogels. Tissue Eng. Part A 00, (2026).

34. Goethe, E. A., Deneen, B., Noebels, J. & Rao, G. The Role of Hyperexcitability in Gliomagenesis. Int. J. Mol. Sci. 24, 1–12 (2023).

35. Naba, A. Mechanisms of Assembly and Remodelling of the Extracellular Matrix. Nature Reviews Molecular Cell Biology vol. 25 (2024).

36. Soko, G. F. et al. Extracellular matrix re-normalization to improve cold tumor penetration by oncolytic viruses. Front. Immunol. 15, 1535647 (2024).

37. Sette, P. et al. GBM-Targeted oHSV Armed with Matrix Metalloproteinase 9 Enhances Anti-tumor Activity and Animal Survival. Mol. Ther. Oncolytics 15, 214–222 (2019).

38. Gao, Y. F. et al. COL3A1 and SNAP91: Novel glioblastoma markers with diagnostic and prognostic value. Oncotarget 7, 70494–70503 (2016).

39. McManis, A. et al. Understanding Glioblastoma Dynamics Using 3D Organoids and Engineered Extracellular Matrix. Advanced Science https://doi.org/10.1002/advs.202522926 (2026) doi:10.1002/advs.202522926.

40. Mohiuddin, E. & Wakimoto, H. Extracellular matrix in glioblastoma: opportunities for emerging therapeutic approaches. Am. J. Cancer Res. 11, 3742–3754 (2021).

41. Yang, Y. et al. Reovirus resistance in tumors mediated by elevated ISG expression : overcoming therapeutic resistance via JAK / STAT pathway modulation. (2026).

42. Qin, Y. et al. Targeting leucine-rich repeat kinase 2 overcomes resistance to oncolytic herpes simplex virus-based therapies in glioblastoma. (2026).

43. Kurokawa, C. et al. Constitutive Interferon Pathway Activation in Tumors as an Efficacy Determinant Following Oncolytic Virotherapy. 110, 1123–1132 (2018).

44. Larrieux, A. & Sajuan, R. Cellular resistance to an oncolytic virus is driven by chronic activation of innate immunity. iScience https://doi.org/10.1016/j.isci.2022.105749 (2023) doi:10.1016/j.isci.2022.105749.

45. Gesundheit, B. et al. Effective Treatment of Glioblastoma Multiforme With Oncolytic Virotherapy: A Case-Series. Front. Oncol. 10, 1–6 (2020).

46. da Hora, C. C., Schweiger, M. W., Wurdinger, T. & Tannous, B. A. Patient-Derived Glioma Models: From Patients to Dish to Animals. Cells 8, (2019).

