## Supplemental material for "Unravelling resistance mechanisms of oncolytic viruses in glioblastoma"

### Supplemental Table

***Table S1: Normalized enrichment scores and p-values of the gene set ‘Neurogenesis’ in all parental-resistant population comparisons.***

| | CME016 – H1PV | LBT123 – H1PV | LBT123 – SINV | LBT007i – SNIV |
|---|---|---|---|---|
| **Normalized enrichment score** | -1.13 | -1.39 | -1.15 | -1.78 |
| **p-value** | 0.013 | $1.3x10^{-7}$ | 0.013 | $4.9x10^{-25}$ |

### Supplemental Figures

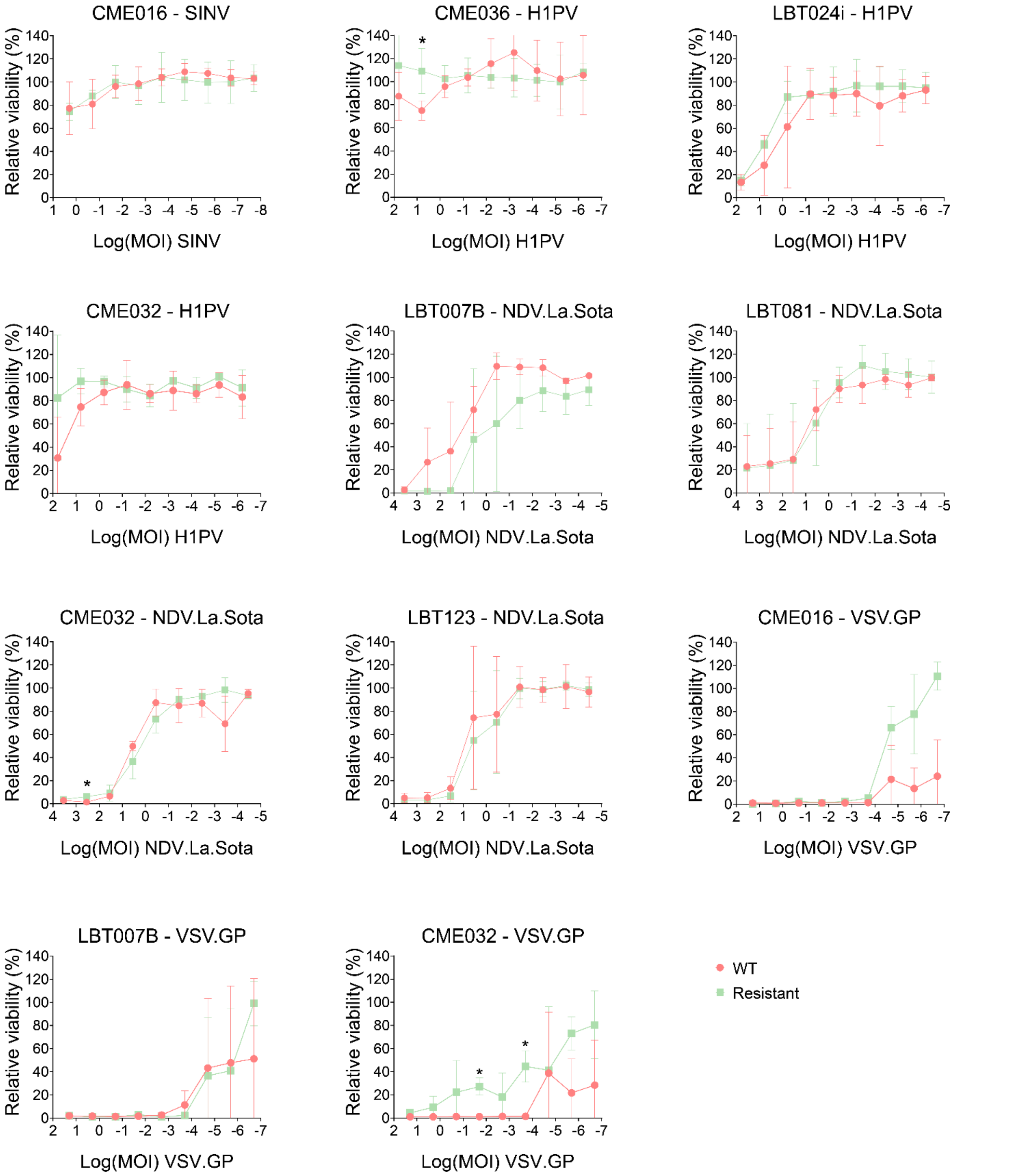


***Figure S1: Overview of non-resistant glioblastoma patient-derived cell line after selection with an oncolytic virus.*** *Viability, relative to uninfected control is shown on the y-axis, whereas the logarithmic function of the multiplicity of infection (MOI) of the virus is shown on the x-axis. An unpaired t test was used to compare the cell viabilities per MOI between resistant and wild type (WT) GBM-PDCLs. Mean of 2 experiments is shown and standard deviations are indicated. *=p<0.05.*

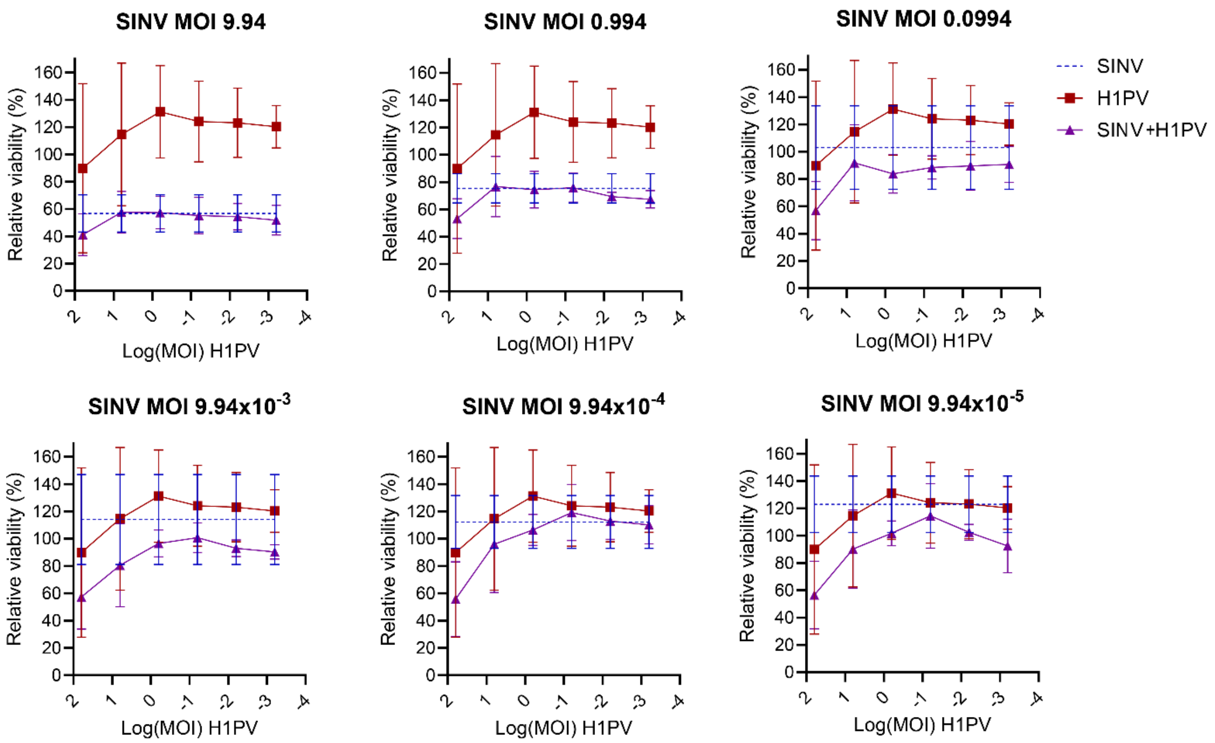


***Figure S2: Simultaneous infection of the glioblastoma patient-derived cell line CME016 with Sindbis and H1-parvovirus.***
*CME016 cells were infected with six 10-fold serial dilutions of Sindbis (SINV) and H1-parvovirus (H1PV). Each SINV multiplicity of infection (MOI) was paired with each H1PV MOI. Cell viability, expressed relative to the uninfected control, is shown on the y-axis, and the log10-transformed H1PV MOI is shown on the x-axis. Each panel represents a single SINV MOI (indicated above the graph) and shows the effects of H1PV alone and in combination with that SINV MOI. Data represent the mean of three independent experiments and standard deviations are indicated. Two-way ANOVA followed by Bonferroni-corrected multiple comparisons revealed no significant decrease in relative viability for any SINV–H1PV combination compared with the corresponding SINV-only treatment.*

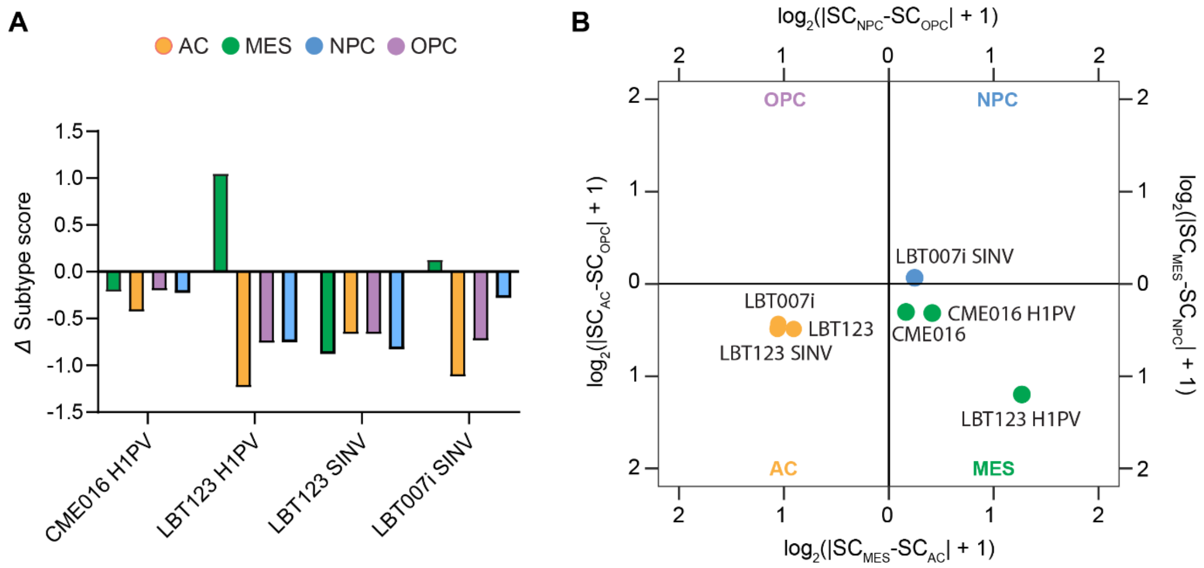


***Figure S3: Neftel subtype score shifts in Sindbis- and H1-parvovirus-resistant versus wildtype patient-derived glioblastoma cell lines.***
***(A)*** *Change in Neftel subtype score calculated in both Sindbis (SINV)- and H1-parvovirus (H1PV)-resistant GBM-PDCLs by subtracting the wildtype (WT) GBM-PDCL subtype score from that of the corresponding resistant glioblastoma patient-derived cell line (GBM-PDCL).* ***(B)*** *Subtype classification of both SINV and H1PV-resistant GBM-PDCLs and their corresponding WT parental lines.*

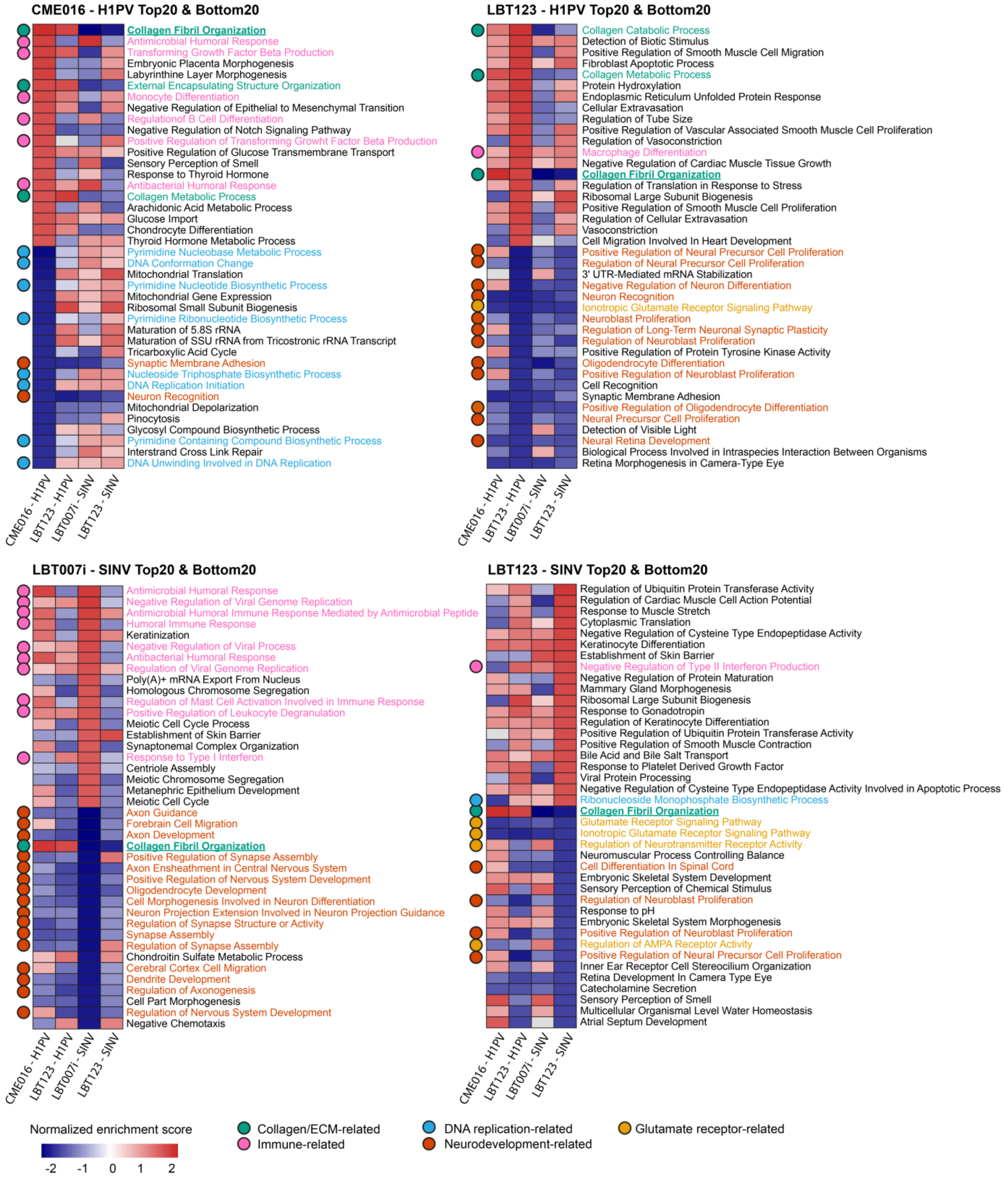


***Figure S4: Gene sets associated with H1-parvovirus and Sindbis virus resistance per glioblastoma patient-derived cell line.***

*Differential gene expression analysis between each H1-parvovirus (H1PV)- and Sindbis virus (SINV)-resistant glioblastoma patient-derived cell lines (GBM-PDCLs) and their WT counterpart was followed by gene set enrichment analysis. The 20 gene sets with the highest (top) and lowest (bottom) normalized enrichment scores were selected for each GBM-PDCL. Gene sets related to collagen/extracellular matrix organization (green), immune response (pink), DNA replication (blue), neurodevelopment (dark orange) and glutamate receptor signaling (yellow) are indicated. Gene sets overlapping in all four GBM-PDCLs are indicated in bold and are underlined.*

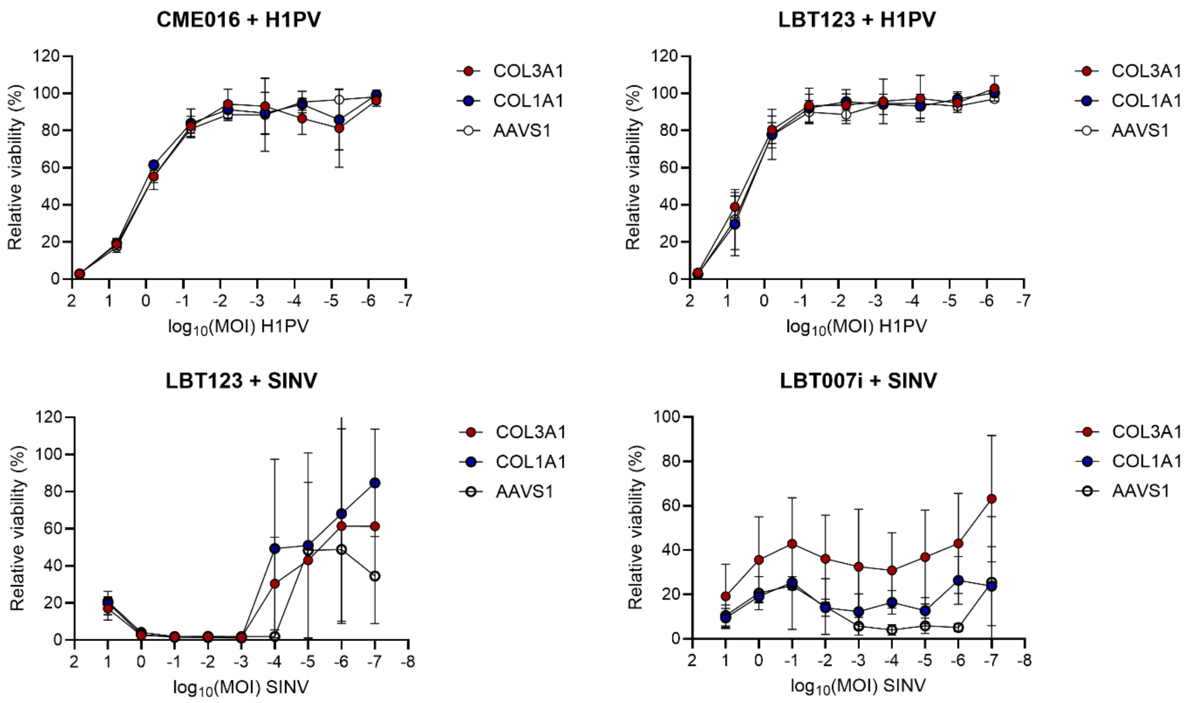


***Figure S5: Knock-out of driver genes COL3A1 and COL1A1 do not influence sensitivity to H1-parvovirus or Sindbis virus.***

*Viability, relative to uninfected control is shown on the y-axis, whereas the logarithmic function of the multiplicity of infection (MOI) of the virus is shown on the x-axis. Driver genes associated with "Collagen fibril organization", COL3A1 and COL1A1, were knocked-out via delivery of viral-like particles in the corresponding parental line of each H1-parvovirus or Sindbis-resistant glioblastoma patient-derived cell line. Viability following infection was compared to an AAVS1 control. Standard deviations are indicated. A paired t test was performed between COL3A1 vs AAVS1 and COL1A1 vs AAVS1 for each MOI. Mean of 2 experiments is shown and standard deviations are indicated.*
